# HIV-1 Unspliced RNA Expression Induces Innate Immune Activation in Macrophages

**DOI:** 10.1101/174508

**Authors:** Hisashi Akiyama, Caitlin M. Miller, Chelsea R. Ettinger, Anna C. Belkina, Jennifer Snyder-Cappione, Suryaram Gummuluru

**Author notes:** Corresponding author: Suryaram Gummuluru, Ph.D.Department of Microbiology Boston University School of Medicine 72 E. Concord St., R512 Boston, MA02118.

## Abstract

Low-levels of type I interferons (IFN-I) are thought to be a driving force for immune activation and T cell exhaustion in HIV-1 infected individuals on highly active antiretroviral therapy (HAART), though the causative mechanisms for persistent IFN-I signaling have remained unclear. Here, we show Rev-CRM1 dependent nuclear export and peripheral membrane association of intron-containing HIV-1 unspliced RNA (usRNA), independent of primary viral sequence or viral protein expression, is subject to sensing, and results in IFN-I-dependent pro-inflammatory responses in macrophages. Our findings suggest that persistent expression of HIV-1 usRNA in macrophages contributes to chronic immune activation and that use of HIV RNA expression inhibitors as adjunct therapy might abrogate aberrant inflammation and restore immune function in HIV-infected individuals on HAART.

## INTRODUCTION

A hallmark of HIV-1 infection in vivo is systemic chronic immune activation (*1*), which has been postulated to lead to HIV-associated non-AIDS complications (HANA) (*2*) and dysfunction of T cells (*3*). Despite long-term viral suppression by HAART and restoration of CD4+ T cell levels, immune activation and inflammation persist in the majority of treated HIV-infected individuals, and is associated with excess risk of mortality and morbidity. Many factors have been attributed to cause this aberrant immune activation in vivo, such as bacterial endotoxin or co-infections (*4*); however, a viral (HIV) etiology for the chronic inflammatory state has remained unclear. Persistent infection of myeloid cells, most likely tissue-resident macrophages, is postulated to contribute to chronic immune activation and HANAs (*5-7*), though molecular mechanisms of how HIV-1 replication activates macrophages remain poorly understood.

In this study, we report that expression and Rev–CRM1 dependent nuclear export of HIV-1 unspliced RNA (usRNA) activates host sensing mechanisms and production of type I interferon (IFN-I)-dependent pro-inflammatory responses in productively infected macrophages. Ability of cells to distinguish intron-containing HIV-1 usRNA from self mRNA was dependent on localization of non-self HIV usRNA at peripheral membrane sites. These findings suggest that novel therapeutic strategies that suppress viral usRNA expression and IFN-I signaling cascades in tissue macrophages might have immunologic and therapeutic benefit in HIV-1 infected individuals on HAART.

## RESULTS

### Late step of HIV-1 replication in macrophages triggers immune activation

HIV-1 infection of monocyte-derived macrophages (MDMs) results in induction of a myeloid cell specific ISG, CD169/Siglec1 (**Fig 1A**) (*8*) whose expression is dramatically up-regulated (5-fold) even upon low levels (<0.3 U/ml) of IFN-α exposure (data not shown). Pre-treatment of MDMs with HIV-1 fusion (maraviroc), reverse transcription (AZT) or integration (raltegravir) inhibitors that blocked establishment of virus infection (**Fig 1B**), abrogated induction of ISGs, CD169 (**Fig 1C**) and IP-10 (CXCL10) (**Fig 1D**). Interestingly, inhibition of Tat-dependent transcription (flavopiridol), but not virus particle maturation (treatment with HIV-1 protease inhibitor, indinavir), in HIV-infected MDMs (**Fig 1B**), blocked enhancement of CD169 expression (**Fig 1C**) and secretion of IP-10 (**Fig 1D**), suggesting that a post-transcriptional step in HIV-1 replication cycle activates MDMs. Furthermore, induction of IFN-β mRNA expression in productively infected MDMs was detected only at 3 days post infection (**Fig 1E**), which was coincident with the up-regulation of CD169 (data not shown), further supporting the hypothesis that a late event in virus replication cycle induces IFN-I responses. Moreover, B18R, IFN-I neutralizing reagent, potently inhibited CD169 expression on MDMs (**Fig 1G**) and reduced IP-10 secretion (**Fig 1H**), confirming the presence of low levels of bioactive IFN-I (below the detection limit of a highly sensitive bioassay (**Fig 1I**)) in HIV-infected MDM culture supernatants that had negligible impact on virus infection (spread) (**Fig 1F**). Collectively, these results suggest that host sensing of a late step of HIV-1 replication in MDMs induces IFN-I-dependent pro-inflammatory responses.

**Fig 1.**
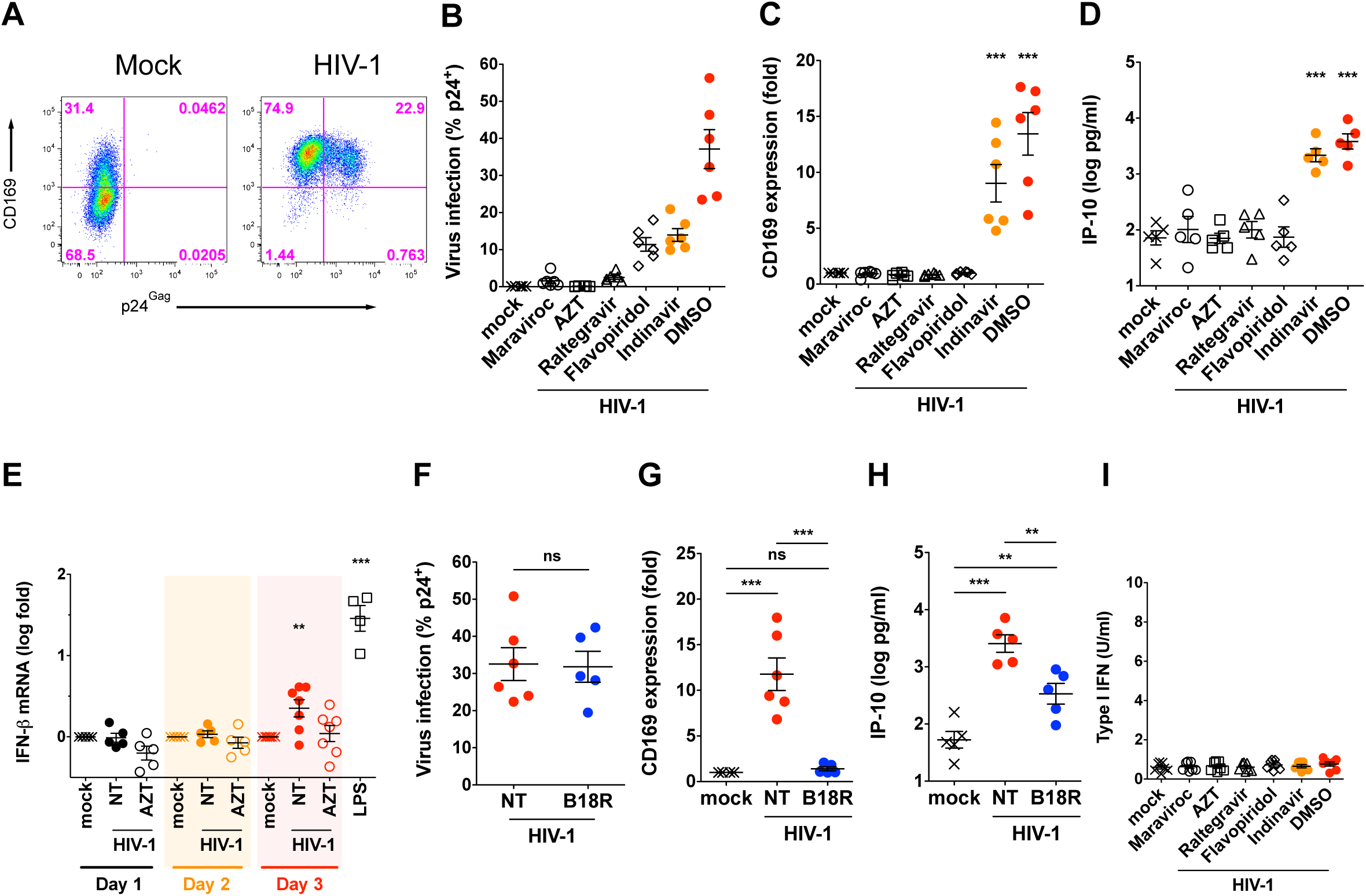
Late step of HIV-1 replication in macrophages triggers immune activation. (A) Representative flow cytometry profiles of HIV-1-infected MDMs analyzed for CD169 expression and HIV-1 infection. (B) HIV-1 infection in MDMs in the presence or absence of inhibitors was measured by intracellular p24^Gag^ staining. (C) CD169 expression (normalized to mock) and (D) IP-10 production in HIV-1-infected MDMs cultured in the presence of anti-HIV-1 drugs. (E) IFN-β mRNA (normalized to untreated) expression in MDMs was quantified. (F) HIV-1 replication (intracellular p24^Gag^), (G) CD169 expression (normalized to mock) and (H) IP-10 production in HIV-1-infected MDMs cultured in the presence B18R. NT: untreated (DMSO). (I) IFN-I secretion from HIV-1-infected MDMs was measured by a bioassay with MDM supernatants harvested on 6 days post infection. The means ± SEM are shown and each symbol represents an independent experiment. Two-tailed p values: one-way ANOVA or paired t-test (F), **: p ≤ 0.01, ***: p ≤ 0.001, ns: not significant.

### Rev–CRM1 dependent HIV-1 usRNA export is subject to host sensing mechanisms

Though induction of pro-inflammatory responses have been described upon exposure to HIV-1 accessory or structural proteins in diverse cell types (*9, 10*), MDMs infected with Nef, Env, Vpu, Vpr, or Vif-deficient mutants (**Fig 2A**) displayed robust induction of CD169 and IP-10 expression (**Fig 2B and C)**. Moreover, MDM infections with Gag p6 (Δp6), NC (ΔNCp6), MA (ΔMA) truncation mutants, or Gag start codon mutant (ATG*) (**Fig 2D, E and H**) that results in initiation of Gag translation from an in-frame internal ATG present at the N-terminus of CA (leading to expression of aberrant Gag) failed to abrogate CD169 or IP-10 induction (**Fig 2F, G, I and J**). Additionally, infection of MDMs with HIV-1 budding deficient mutant (PTAP-) (*11*), CA-mutants deficient for Gag–Gag interaction (*12*), and cyclophilin A (CyPA)-binding-deficient CA mutant G89V (**Fig 2K, N and Q**) resulted in up-regulation of CD169 and IP-10 (**Fig 2L, M, O, P, R and S**), suggesting that neither virion-release (*13*), cytoplasmic accumulation of higher-ordered Gag assembly intermediates nor CyPA-binding to de novo expressed Gag, a target of a “cryptic sensor” in myeloid dendritic cells (*10*), is required for MDM activation. Furthermore, complete abrogation of Gag-pol expression (ΔGag-pol; **Fig 2D**) only modestly attenuated expression of CD169 and IP-10 (**Fig 2U and V**) in infected MDMs (**Fig 2T**), suggesting that structural and accessory proteins of HIV-1 do not encode immune-activation determinants.

**Fig 2.**
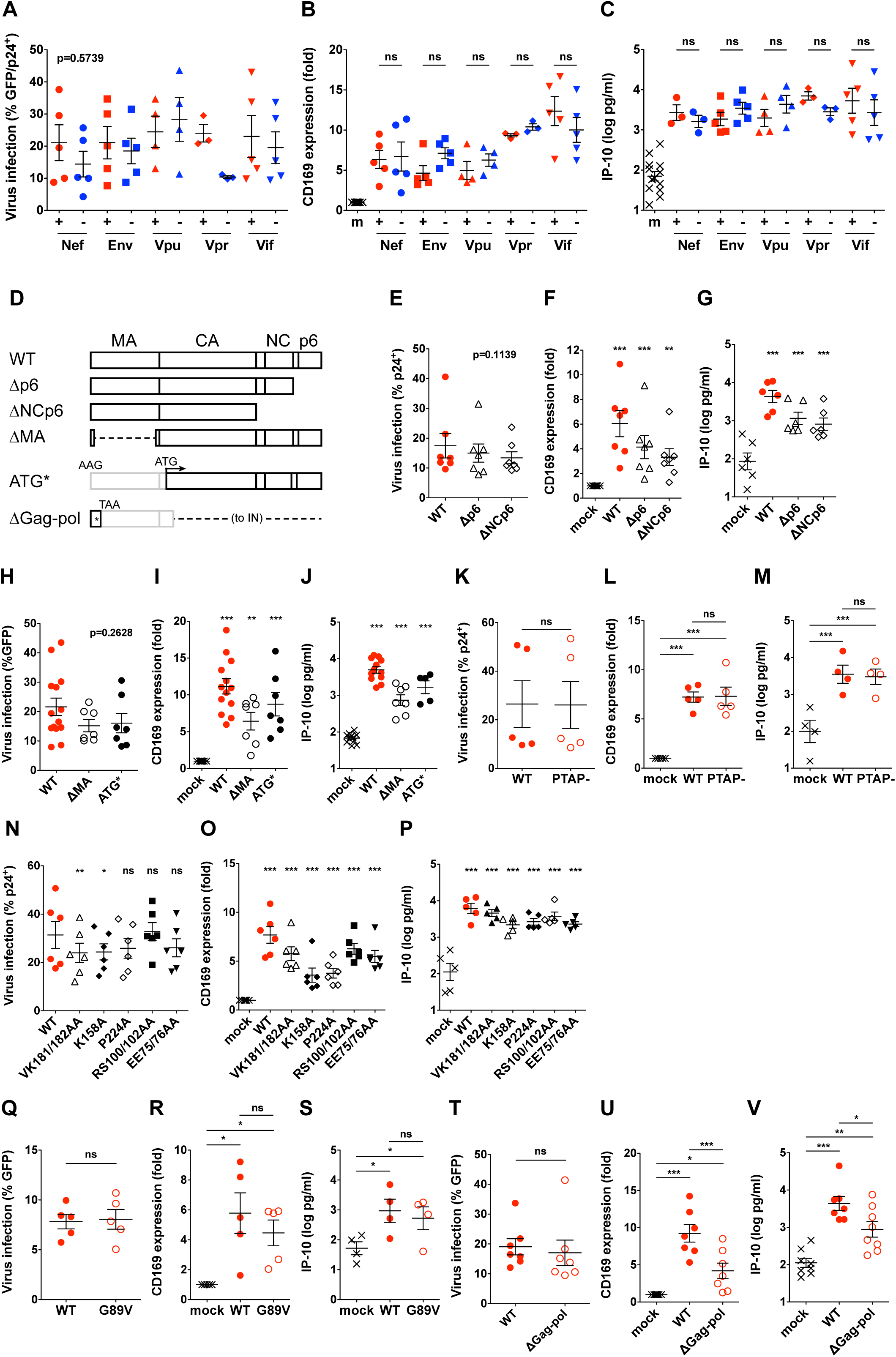
Structural and accessory proteins of HIV-1 do not encode immune-activation determinants. (A) HIV-1 infection (GFP expression or intracellular p24^Gag^), (B) CD169 expression (normalized to mock), and (C) IP-10 production in MDMs infected with indicated HIV-1 mutants. m: mock. (D) Schematic demonstrations of HIV-1 Gag mutants. (E) HIV-1 infection (intracellular p24^Gag^), (F) CD169 expression (normalized to mock) and (G) IP-10 production in MDMs infected with HIV-1 mutants lacking p6 or NC-p6. (H) HIV-1 infection (GFP expression), (I) CD169 expression (normalized to mock), and (J) IP-10 production in MDMs infected with HIV-1 mutants lacking MA. (K) HIV-1 expression (intracellular p24^Gag^), (L) CD169 expression (normalized to mock), and (M) IP-10 production in MDMs transduced with the wild type (WT) or PTAP-mutant. (N) HIV-1 expression (intracellular p24^Gag^), (O) CD169 expression (normalized to mock), and (P) IP-10 production in MDMs transduced with the wild type (WT) or indicated CA mutants deficient for intra- or inter-hexamer formation. (Q) HIV-1 infection (GFP expression), (R) CD169 expression (normalized to mock), and (S) IP-10 production in MDMs transduced with the wild type (WT) or CyPA-binding-deficient mutant (G89V). (T) HIV-1 infection (GFP expression), (U) CD169 expression (normalized to mock), and (V) IP-10 production in MDMs infected with HIV-1 ΔGag-pol mutant. The means ± SEM are shown and each symbol represents an independent experiment. Two-tailed p values: one-way ANOVA or paired t-test (K, Q and T), *: p ≤ 0.05, **: p ≤ 0.01, ***: p ≤ 0.001, ns: not significant.

To identify the post-transcriptional viral product that induces pro-inflammatory responses in MDM, we next assessed the effect of viral RNA on MDM activation. There are three classes of viral mRNAs (multiply-spliced, singly-spliced and unspliced) transcribed from the HIV-1 LTR (*14*). Nuclear export of HIV-1 usRNA relies on Rev– RRE binding and a cellular mRNA transporter CRM1 (*15, 16*). Mutations that disrupt Rev binding to CRM1 (M10 mutant; **Fig 3A**) or delete RRE (ΔRRE; **Fig 3A**) abrogate usRNA nuclear export and thus Gag expression, but not export and translation of multiply-spliced viral RNAs (i.e. GFP in place of *nef*; **Fig 3A, B and C**) (*15, 16*). Interestingly, infection of MDMs with the M10 mutant failed to up-regulate CD169 and IP-10 expression (**Fig 3D and E**). Insertion of the constitutive transporting element (CTE) from Mason-Pfizer monkey virus (MPMV) (*17*) within the *pol* open reading frame (orf) of the M10 mutant (M10-CTE) or the ΔRRE mutant (ΔRRE-CTE) in the sense orientation, but not in the anti-sense orientation (M10-CTE-AS), rescued nuclear export of HIV-1 usRNA and Gag expression (**Fig 3A and B**). However, rescue of HIV-1 usRNA nuclear export by CTE-dependent pathway failed to induce CD169 or IP-10 expression (**Fig 3D and E**). Moreover, while Rev-dependent expression of a non-viral protein (TagRFP) from the intron-containing viral usRNA (**Fig 3F and G**) induced CD169 and IP-10 expression (**Fig 3H and I**), Rev-independent expression of intron-less non-viral protein (ZsGreen) (**Fig 3F and J**) failed to induce CD169 and IP-10 expression (**Fig 3K and L**). These findings suggest that engagement of divergent RNA nuclear export pathways such as those mediated by Rev–RRE and MPMV CTE might lead to altered exposure of viral usRNA to host cytoplasmic sensing machinery, and that host sensing of viral usRNA specifically exported via Rev–CRM1 dependent pathway is necessary for inducing pro-inflammatory responses in MDMs.

**Fig 3.**
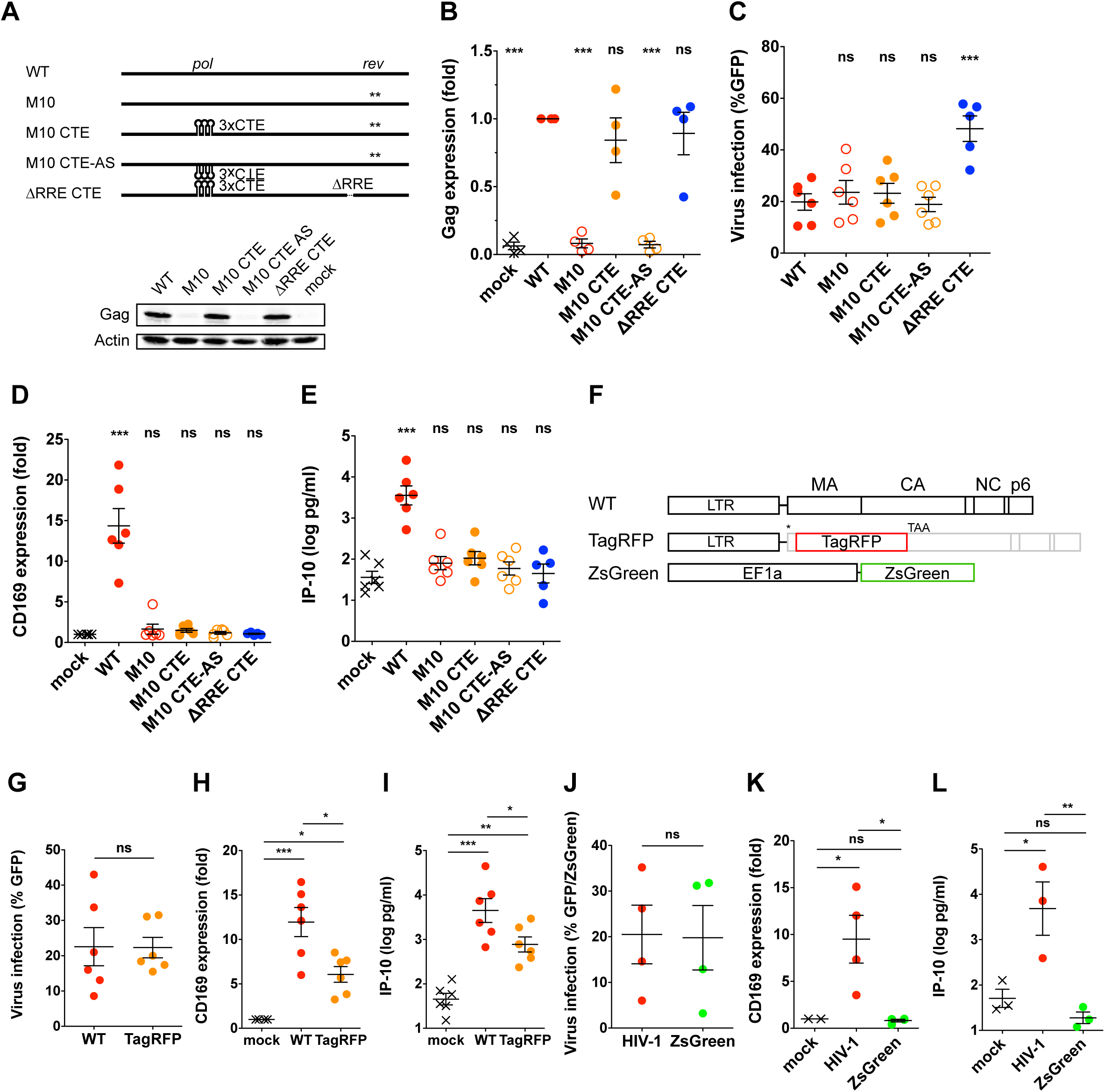
Rev–CRM1 dependent HIV-1 usRNA export is required for sensing. (A) Schematic demonstration of HIV-1 Rev mutants and their Gag expression in infected MDMs. *: position of mutation, CTE: constitutive transport element, RRE: rev responsive element. (B) Quantification of Gag expression in infected MDMs (normalized to WT infection). (C) HIV-1 infection (GFP expression), (D) CD169 expression (normalized to mock), and (E) IP-10 production in MDMs infected with the HIV-1 Rev mutants. (F) Schematic demonstration of TagRFP and ZsGreen expression vectors. (G) HIV-1 infection (GFP expression), (H) CD169 expression (normalized to mock), and (I) IP-10 production in MDMs infected with the TagRFP mutant. (J) Transduction efficiency (GFP or ZsGreen expression), (K) CD169 expression (normalized to mock), and (L) IP-10 production in MDMs transduced with the ZsGreen vector. The means ± SEM are shown and each symbol represents an independent experiment. Two-tailed p values: one-way ANOVA or paired t-test (G, J), *: p ≤ 0.05, **: p ≤ 0.01, ***: p ≤ 0.001, ns: not significant.

### Membrane targeting of HIV-1 usRNA is required for MDM activation

HIV-1 usRNA exported via Rev–RRE or CTE dependent pathway results in distinct cytoplasmic distribution (*18*). We therefore tested the hypothesis that cytoplasmic localization of HIV-1 usRNA-associated ribonucleoprotein complexes (RNPs) specific to the Rev–RRE pathway dictates induction of pro-inflammatory signaling cascades. Since a subset of cytoplasmic HIV-1 usRNAs are trafficked with Gag to membrane-associated virus assembly sites for packaging into new virions (*12, 14*), we infected MDMs with a series of Gag-MA mutants to potentially modulate localization of Gag-containing HIV-1 usRNA-associated RNPs. We found that none of the well-characterized MA membrane targeting domain mutants or those that potentially alter Gag assembly structure or HIV usRNA–Gag association prevented MDM activation (**Fig 4A, B and C**). Membrane-targeting function is well conserved amongst all retroviral MA proteins, though site of virus assembly varies amongst retroviruses (*14*). To putatively alter membrane association sites of HIV-1 usRNA, we next utilized a HIV-1/MLV chimeric virus, mMA12, which contains MA and p12 of MLV in place of HIV-1 MA (**Fig 4D**)(*19*). Interestingly, infection of MDMs with mMA12 attenuated CD169 and IP-10 expression in productively infected MDMs (**Fig 4E, F and G**). Finally, infection of MDMs with myristoylation-deficient MA mutant (G2A) (**Fig 4H**) that prevents stable membrane association of HIV-1 Gag (*12*) attenuated viral usRNA-induced MDM activation (**Fig 4I and J**). These results suggest that stable membrane association of viral usRNA is necessary for host sensing and induction of type I IFN-dependent pro-inflammatory signals in MDMs.

**Fig 4.**
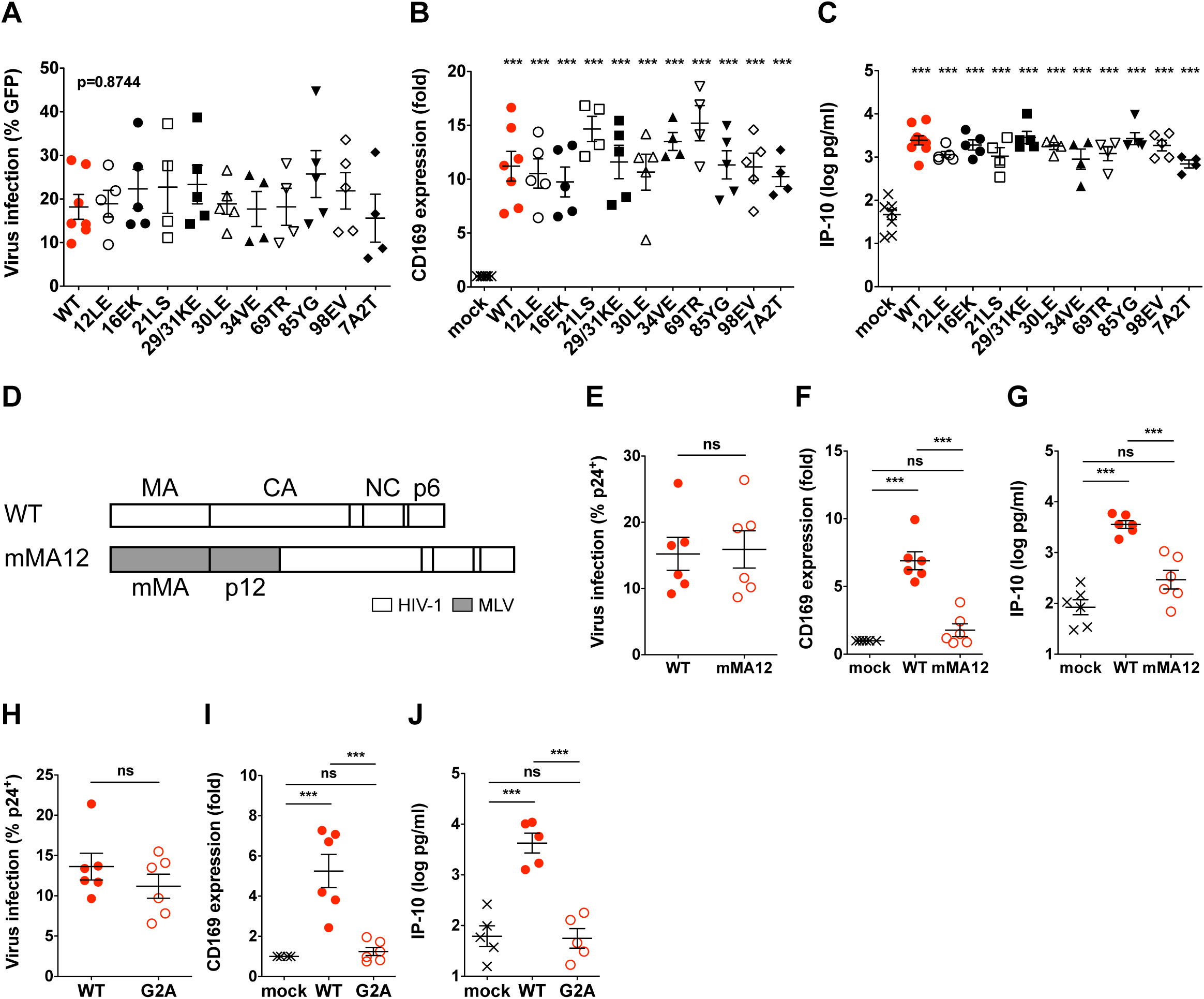
Membrane targeting of viral usRNA is required for MDM activation. (A) Virus infection (GFP expression), (B) CD169 expression normalized to mock, and (C) IP-10 production in MDMs transduced with WT or indicated MA mutants lacking various MA functions (Table 1). (D) Schematic demonstration of MLV/HIV-1 chimera, mMA12. (E) Viral infection (intracellular p24^Gag^), (F) normalized CD169 expression, and (G) IP-10 production in MDMs infected with WT or mMA12 viruses. (H) HIV-1 infection (intracellular p24^Gag^), (I) CD169 expression normalized to mock, and (J) IP-10 production in MDMs infected with WT or G2A mutant. The means ± SEM are shown and each symbol represents an independent experiment. Two-tailed p values: one-way ANOVA or paired t-test (E, H), ***: p ≤ 0.001, ns: not significant.

## DISCUSSION

Whether HIV-1 infection of MDMs is subject to sensing has remained a matter of controversy. In this study, we show that a late step of HIV-1 replication, specifically, expression and Rev–CRM1 mediated nuclear export of viral usRNA and its peripheral membrane association in MDMs is subject to host sensing mechanisms and results in innate immune activation and induction of pro-inflammatory responses. Importantly, we hypothesize that Rev–CRM1 dependent nuclear export specifically “marks” HIV-1 usRNA for detection by cytoplasmic sensing mechanisms, since access to alternative CTE-dependent nuclear export pathway failed to induce pro-inflammatory responses. There seems to be no sequence specificity for the sensing mechanism since inclusion of non-viral sequences (TagRFP) in viral usRNA failed to abrogate production of pro-inflammatory responses. Alternatively, “tethering” of HIV-1 usRNA to distinct peripheral membrane sites appeared to be a critical determinant; altering membrane targeting (by MLV-MAp12) or preventing membrane association (HIV-1 MA-G2A mutant) of viral usRNA potently attenuated induction of pro-inflammatory responses. While the distinct trafficking pattern and composition of viral usRNA-containing RNPs has been hypothesized to confer translational advantage to HIV-1 mRNAs compared to bulk host mRNA (*20*), such differences might also subject viral usRNA to detection by host innate immune sensing mechanisms. We hypothesize that a yet-to-be-identified sensor that resides at or in the vicinity of HIV-1 Gag assembly site senses HIV-1 usRNA-containing RNP complexes and activates MDMs. Further study is warranted to identify viral RNP complexes and the host factor(s) that activate MDMs upon HIV-1 infection.

Tissue-resident macrophages are important cellular targets of HIV, estimated to compromise up to 4% of infected cells *in vivo* (*21*), and can remain persistently infected with HIV-1 even in individuals on HAART (*5-7*). HIV-1 usRNAs have been detected in CD4+ T cells and alveolar macrophages from patients on long-term HAART (*22, 23*). While these RNAs may not lead to functional viral protein production, persistent expression of viral usRNA might trigger host sensing in MDMs, that we demonstrate is a potent stimulus of IFN-I dependent pro-inflammatory responses. It is possible that HIV-1 infection of tissue-resident macrophages induces low levels of IFN-I and bystander myeloid cell activation contributing to systemic immune activation. Unfortunately, current HAART regimen does not block virus RNA transcription in previously infected cells. Thus, the development and clinical application of inhibitors that decrease viral RNA expression such as Tat and Rev inhibitors (*24, 25*), especially in tissue macrophages, could alleviate systemic immune activation in HIV-infected individuals on HAART.

## MATERIALS AND METHODS

### Plasmids

HIV-1 replication competent molecular clones, Lai and Lai/YU-2env, single-round reporter constructs, LaiΔenv and LaiΔenvGFP (GFP in place of the *nef* orf) have been described previously (*26, 27*). HIV-1 mutants, LaiΔenvΔvpr, Lai/YU-2env and Lai/YU-2envΔvpu, have been previously reported (*26, 28*). To create LaiΔenvGFPΔvif, a *vif*-containing fragment from LaiΔenv-lucΔvif (frame-shift insertions at the NdeI site in the *vif* orf) (*29*) was replaced into the corresponding region of LaiΔenvGFP. LaiΔenvGFPΔGag-pol was created by inserting frame-shift mutations at the ClaI site (nt 377) in the Env-expression construct which has a large deletion between N-terminus of CA and IN (*26*) and the fragment containing the *env* and *nef* orfs was replaced with the corresponding portion of LaiΔenvGFP. The CA G89V R9-GFP mutant and its parental virus R9-GFP were generous gifts from Dr. Christopher R. Aiken (Vanderbilt University School of Medicine). The PTAP-mutant and its parental virus R9 were generously provided by Dr. Wesley I. Sundquist (University of Utah Health). The following Gag mutants were generous gifts from Dr. Jaisri Lingappa (University of Washington): VK181/182AA, K158A, P224A, RS100/102AA, EE75/76AA, Δp6, ΔNCp6, G2A. mMA12ΔenvGFP was created by replacing the luciferase gene of MHIV mMA12 (a generous gift from Dr. Masahiro Yamashita, Aaron Diamond AIDS Research Center) with the GFP gene. Rev-deficient mutant (M10) was created by PCR-directed mutagenesis using the primer set (**Table 1**). To make RRE-deletion mutant (ΔRRE), a StuI-AleI fragment in the *env* orf was deleted from LaiΔenvGFP. The CTE was PCR-amplified using the primers (**Table 1**) and pCR4-CTE (a gift from Dr. Mario Chin, Addgene plasmid # 36868) as template and three repeats of CTE were inserted into M10 or ΔRRE between the BclI (nt 2011) and NheI (nt 3467) sites in a sense or antisense direction. HIV-1 MA-deficient mutants, ΔMA (lacking amino acid residues 6 to 125) and ATG*, and the following Gag mutants were created by PCR mutagenesis (QuikChangeII, Agilent) or overlapping PCR using the primers listed in **Table 1**: ΔMA, 12LE, 16EK, 21LS, 29/31KE, 30LE, 34VE, 69TR, 85YG, 98EV, 7A2T using LaiΔenvGFP as a backbone. To express TagRFP in place of HIV-1 MA, an NcoI site was PCR-generated into at nt 378 (41 nt downstream of Gag starting codon) of the ATG* mutant. This vector (HIV-vec-plus) has a potion of HIV-1 genome in order to enhance chimeric viral RNA incorporation into virions (*30*). Then, an AatII site was PCR-generated just before the HIV-1 PR cleavage site into HIV-vec-plus (HIV-vec-plus-AatII) and PCR-amplified TagRFP (Evrogen) was inserted into HIV-1-vec-plus-AatII. A lentivector expressing ZsGreen, pHAGE-ZsGreen, is a generous gift from Dr. Darrell Kotton (Boston University), and HIV-1 packaging plasmid psPAX2 and VSV-G expression constructs have been previously described (*31*).

**Table 1.**
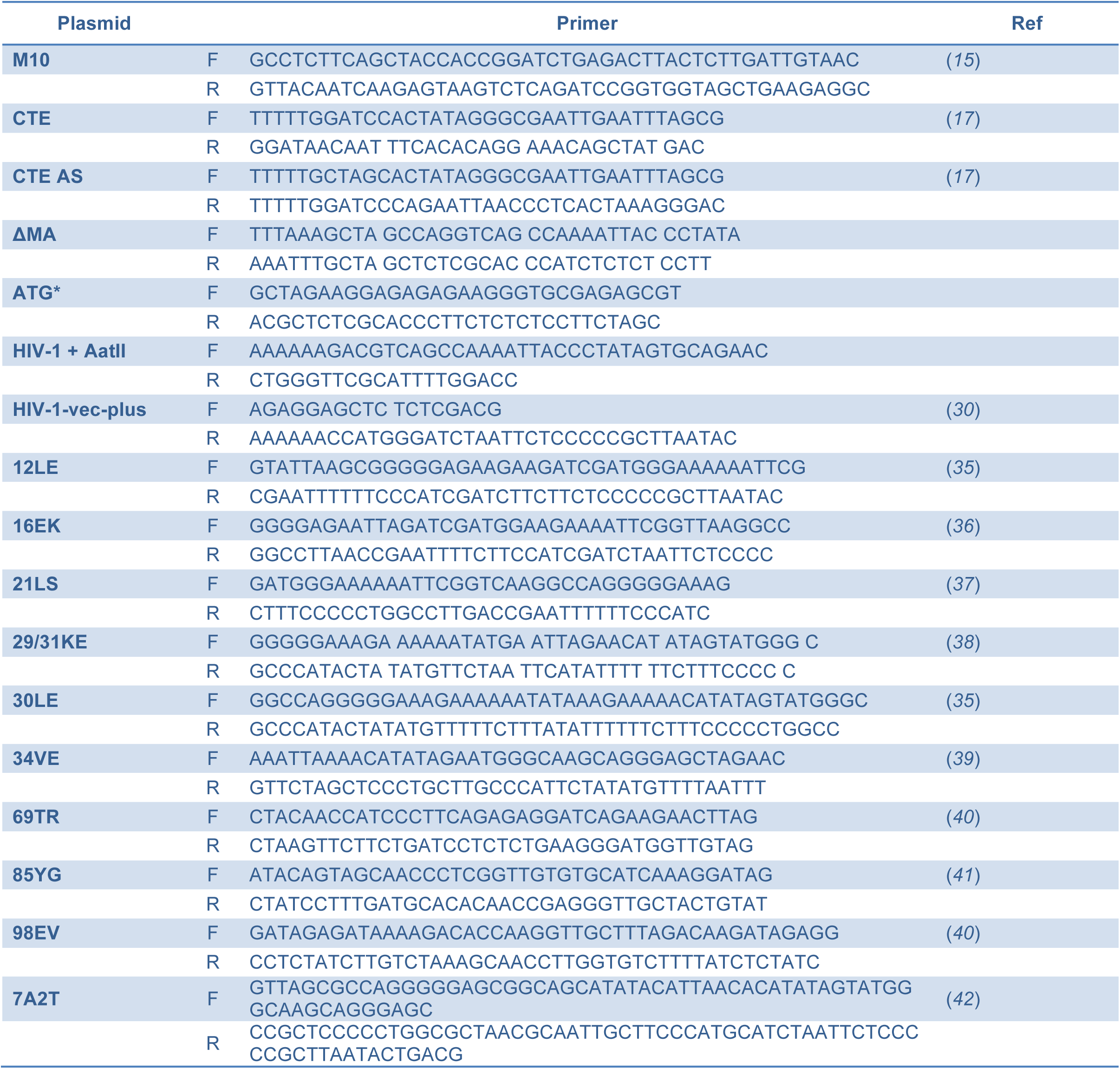

### Cells

Human monocyte-derived macrophages (MDMs) were derived from beads-isolated CD14+ peripheral blood monocytes (*26*) by culturing in RPMI1640 (Invitrogen) containing 10% heat-inactivated human AB serum (Gemini Bio Products, Sigma or Corning) and recombinant human M-CSF (20 ng/ml; Peprotech) for 5-6 days. HEK293T, 293 ISRE-luc and TZM-bl have been described previously (*28, 31, 32*).

### Viruses

Replication competent viruses were derived from HEK293T cells via calcium phosphate transient transfection of HEK293T cells by calcium phosphate as described previously (*31*). Single-round-replication-competent viruses pseudotyped with VSV-G were generated from HEK293T cells via co-transfection of HIV-1Δenv constructs and VSV-G expression plasmid. To express HIV-1 mutants which by themselves cannot infect MDMs, a mutant plasmid was co-transfected with the packaging construct (psPAX2) and VSV-G expression vector into HEK293T (*31*). Virus-containing cell supernatants were harvested 2 days post-transfection, cleared of cell debris by centrifugation (300 x g, 5 min), passed through 0.45 µm filters, and purified and concentrated by ultracentrifugation on a 20% sucrose cushion (24,000 rpm and 4°C for 2 hours with a SW32Ti or SW28 rotor (Beckman Coulter)). The virus pellets were resuspended in PBS, aliquoted and stored at -80 °C until use. The capsid content of HIV-1 was determined by a p24gag ELISA (*31*) and virus titer was measured on TZM-bl (*33*).

### Infection

MDMs seeded one day before infection were spinoculated with viruses (1h at RT and 2,300 rpm) in the presence of polybrane (Milipore) at various moi (typically 0.5 to 2), cultured for 2-3 hours at 37°C, washed to remove unbound virus particles, and cultured for 3-6 days. Infection in MDMs was quantified by analyzing intracellular p24^Gag^ or GFP expression by flow cytometry. In some experiments, MDMs were pretreated prior to infection or treated post infection with the following drugs/reagents: maraviroc (NIH AIDS Reagent Program), AZT (NIH AIDS Reagent Program), raltegravir (NIH AIDS Reagent Program or Selleckchem), flavopiridol (NIH AIDS Reagent Program), indinavir (NIH AIDS Reagent Program) and B18R (eBioscience).

### Immunoblot analysis

To assess expression of p24gag, cell lysates containing 20-30 µg total protein were separated by SDS-PAGE, transferred to nitrocellulose membranes and the membranes were probed with a mouse anti-p24 antibody (p24-2, Dr. Michael H. Malim, NIH AIDS Reagent Program) followed by donkey anti-goat-IgG-IRDye 680 (Pierce). As loading controls, actin was probed using rabbit anti-actin (SIGMA) followed by a goat anti-rabbit-IgG-IRDye 800CW (Pierce). Membranes were scanned with an Odessy scanner (Li-Cor).

### mRNA quantification

Total mRNA was isolated from 1x106 cells using a kit (RNeasy kit, QIAGEN) and reverse-transcribed using oligo(dT)20 primer (SuperscriptIII, Invitrogen). IFN-β mRNA was quantified using Maxima SYBR Green (Thermo Scientific) and normalized to GAPDH mRNA by the 2^-ΔΔ*C*^_T_ method as described (*28, 34*). As a positive control for IFN-β mRNA expression, MDMs were treated with 1 µg/ml LPS (InvivoGen) for 2 hours.

### Cytokine quantification

IP-10 in MDM culture supernatants was measured with a BD Human IP-10 ELISA Set (BD). Bioactive IFN-I was measured by a bioassay using 293 ISRE-luc cell line (*28*).

### Flow cytometry

CD169 on MDMs was stained with Alexa647-condjugated mouse anti-human CD169 antibody (BioLegend) and analyzed with BD LSRII (BD). Geo mean fluorescence intensity (MFI) was calculated and normalized to that of uninfected (mock) MDMs. In some experiments, intracellular p24 was stained as described (*28*) using FITC- or RD1-condjucated mouse anti-p24 monoclonal antibody (KC57, Coulter). Data was analyzed with FlowJo software (FlowJo).

### Statistics

All the statistics analysis was performed using GraphPad Prism 5. Two-tailed p values were calculated using one-way ANOVA followed by the Tukey-Kramer post-test (symbols for p values shown with a line) or the Dunnett’s post-test (comparing to control (mock or wild type), symbols for p values shown on each column), or a paired t-test (comparing two samples, symbols for p values shown with a line). Symbols represent, *: p ≤ 0.05, **: p ≤ 0.01, ***: p ≤ 0.001, ns: not significant (p > 0.05).

## Acknowledgments

We thank Dr. Gregory A. Viglianti and Dr. Andrew J. Henderson for discussions and for critical reading of the manuscript. We thank Dr. Wesley I. Sundquist, Dr. Christopher R. Aiken, Dr. Jaisri Lingappa, Dr. Masahiro Yamashita and Dr. Darrell Kotton for generous gifts of reagents. We thank the BUMC Flow Cytometry Core and the Analytical Instrumentation Core for technical assistance. This work was supported by NIH grants AI064099 (SG), HD083111 (SG) and P30AI042853 (SG).

